# On the deformability of an empirical fitness landscape by microbial evolution

**DOI:** 10.1101/293407

**Authors:** Djordje Bajić, Jean C.C. Vila, Zachary D. Blount, Alvaro Sánchez

**Author notes:** These authors contributed equally (Djordje Bajić, Jean C.C. Vila). Coresponding authors: DB and AS.

## Abstract

A fitness landscape is a map between the genotype and its reproductive success in a given environment. The topography of fitness landscapes largely governs adaptive dynamics, constraining evolutionary trajectories and the predictability of evolution. Theory suggests that this topography can be “deformed” by mutations that produce substantial changes to the environment. In spite of its importance, the deformability of fitness landscapes has not been systematically studied beyond abstract models, and little is known about its reach and consequences in empirical systems. Here we have systematically characterized the deformability of the genome-wide metabolic fitness landscape of the bacterium *E. coli*. Deformability is quantified by the non-commutativity of epistatic interactions, which we experimentally demonstrate in mutant strains on the path to an evolutionary innovation. Our analysis shows that the deformation of fitness landscapes by metabolic mutations rarely affects evolutionary trajectories in the short-range. However, mutations with large environmental effects leave these as a “legacy”, producing long-range landscape deformations in distant regions of the genotype space that affect the fitness of later descendants. Our methods and results provide the basis for an integration between adaptive and eco-evolutionary dynamics with complex genetics and genomics.

## Introduction

When a new genotype appears in a population its reproductive success is largely governed by the environment. Although it is often thought of as an external driver of natural selection, the environment can also be shaped by the population itself, for instance through its metabolic activity, or through interactions with the abiotic habitat or other species (F. John Odling-Smee, Kevin N. Laland, Marcus W. Feldman, 2003; Laland et al., 2014; Lewontin, 1983). These population-driven environmental changes can in turn modify the fitness effects of future mutations, closing in an “eco-evolutionary” feedback loop (Post and Palkovacs, 2009). Eco-evolutionary feedbacks are well documented in natural (Hendry, 2016) and experimental populations (Jones et al., 2009), and at all scales of biological organization: from the cellular scale, e.g. in the evolution of cancer (Basanta and Anderson, 2017) and microbial populations (Sanchez and Gore, 2013), to the organismal scale in animal (Matthews et al., 2016) and plant evolution (terHorst and Zee, 2016).

The one-to-one “map” between each genotype and its adaptive value in a given environment is known as the fitness landscape (Wright, 1932). Since populations actively modify their environment, new mutations can in principle have environmental as well as fitness effects. Thus, evolving populations may reshape or “deform” the fitness landscapes on which they are adapting (Kauffman and Johnsen, 1991). Though they are often used only metaphorically to depict or visualize adaptation, fitness landscapes are a major determinant of evolution. In particular, the topography of a fitness landscape (i.e. the location of fitness peaks and valleys and their connectivity) plays a pivotal role, as it governs the accessibility of evolutionary trajectories (Hartl, 2014; Poelwijk et al., 2007; Weinreich et al., 2006); the role of population structure on evolution (Nahum et al., 2015); the degree of evolutionary convergence among populations (Van Cleve and Weissman, 2015); the expected role of drift, selection, and sex in the evolutionary process (Moradigaravand and Engelstädter, 2012; Rozen et al., 2008); the discovery of evolutionary innovations (Barve and Wagner, 2013); and the predictability of evolution (de Visser and Krug, 2014), a subject of growing importance for the management of pathogens and cancer treatment (Barber et al., 2015; Lässig et al., 2017; Luksza and Lässig, 2014; Nourmohammad et al., 2013; Zhao et al., 2016). Given the critical role it plays in adaptation, if populations do indeed change the topography of their fitness landscapes as they evolve, it is imperative to understand precisely how. Do mutations that alter the environment generally deform the fitness of every other subsequent mutation, or just a subset of them? If the latter, where are those “deformations” localized in the genotype space, and how strong are they? All of these questions remain open as the deformability (or “rubberness”) of fitness landscapes has never been systematically studied beyond abstract theoretical models (Kauffman and Johnsen, 1991). Substantial experimental evidence suggests that microbial fitness landscapes are likely to exhibit deformability (Friesen et al., 2004; Gac and Doebeli, 2010; Good et al., 2017; Paquin and Adams, 1983; Rosenzweig et al., 1994), making microbes an ideal system with which to address this issue. Microbial metabolism leads to large-scale environmental construction through the uptake and release of metabolites (Good et al., 2017; Rosenzweig et al., 1994). Which nutrients are uptaken, which byproducts are produced and released, and in what amounts, are all governed by the structure of the metabolic network and, therefore, by the genotype (Paczia et al., 2012; Quandt et al., 2015). Therefore, new mutations that change the metabolic network can also change the patterns of metabolic uptake and secretion, altering the environment and potentially also the fitness of future mutations (Rosenzweig et al., 1994). Microbial physiology and growth can be explicitly simulated using genome-scale metabolic models (Lewis et al., 2010; O’Brien et al., 2015; Orth et al., 2011). Due to their excellent predictive capabilities (Orth et al., 2011) and ability to easily and rapidly screen millions of genotypes, these genome-wide metabolic models have been successfully used to systematically explore the genotype space (Matias Rodrigues and Wagner, 2009). Recent advances in dynamic metabolic modeling make it possible to explicitly simulate the growth of microbial communities and their environmental feedbacks with evolution (Harcombe et al., 2014; Mahadevan et al., 2002), making genome-wide dynamic metabolic modeling of microbial genotypes a promising method to examine the deformability of fitness landscapes (Fig. 1A).

**Fig. 1.**
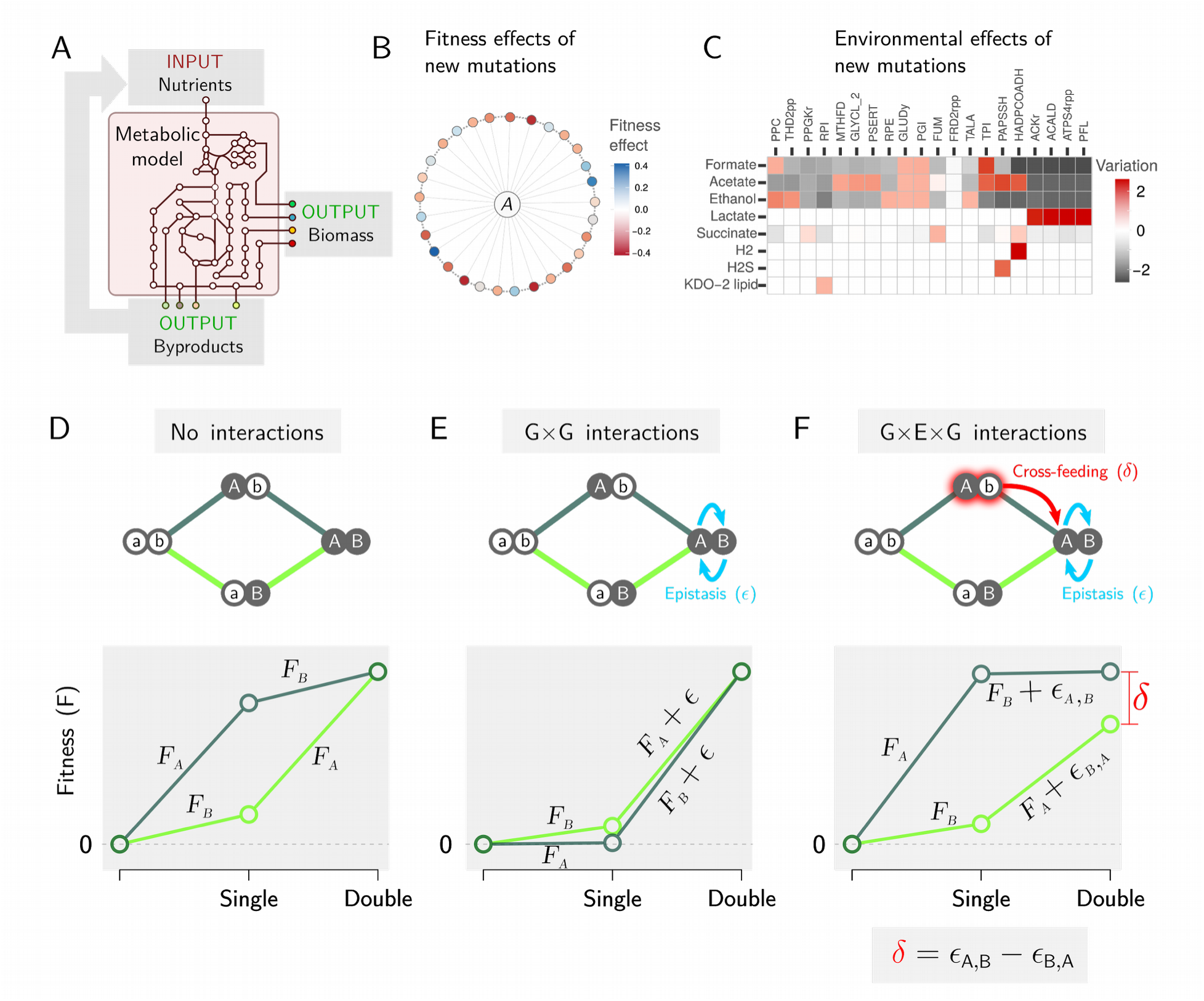
Measuring deformability in the *E. coli* metabolic fitness landscape. **A**. Schematic depiction of dynamic Flux Balance Analysis (dFBA) simulations. Given an input in the form of nutrients, metabolic fluxes through an explicit and empirically curated metabolic model are optimized to maximize the biomass growth yield. The optimal metabolic fluxes produce metabolic byproducts that are released to the external environment, becoming part of it and of future inputs. **B.** A small subset of genotypes differing from our *E. coli* metabolic model in a single mutation (an added or deleted reaction), colored according to its effect on fitness in competition with the ancestor (A). **C.** Environmental effects of a subset of mutants, expressed as the variation in the profile of secreted metabolites compared to ancestral *E. coli* genotype (computed as log-modulus transformed difference in the amount of a given secreted molecule; see Methods). Mutant names given as reaction names in BIGG database notation. **D.** Two loci fitness landscape in the absence of gene-gene interactions, where the fitness of each mutation is the same in both the ancestral and the other mutant’s genetic backgrounds. Fitness of each genotype was calculated in direct competition with its immediate ancestor. Mutations A and B correspond to the addition of GLYCL_2 (in BiGG database notation) glycine cleavage system) and AIRCr (phosphoribosylaminoimidazole carboxylase) respectively. **E.** Two-loci fitness landscape with gene-gene interactions giving rise to epistasis (**ε**). Mutations A and B were respectively SO3R (sulfite reductase) and PAPSSH (phosphoadenylylsulfatase), simulated in a constant environment. **F.** Two-loci fitness landscape where one of the mutants transforms the environment, leading to cross-feeding towards the double mutant. Mutations A and B correspond to the addition of PAPSSH and HADPCOADH (3-hydroxyadipyl-CoA dehydrogenase), simulated in conditions where environmental construction was allowed. In addition to regular epistasis, this leads to a non-commutative epistatic shift (δ).

Here we have used such an approach, as well as experiments with the bacterium *E. coli*, to show that fitness landscape deformability can profoundly alter the definition of epistasis, which becomes dependent on the order in which mutations occur. By systematically screening the *in silico* metabolic fitness landscape of *E. coli*, we are able to offer a precise view of how deformability by eco-evolutionary feedbacks plays out over short and long mutational ranges.

## Results

### Non-commutative epistasis characterizes fitness landscape deformability

To investigate the effect of metabolic secretions on the fitness landscape, we used dynamic Flux Balance Analysis (dFBA) to determine the distribution of fitness and environmental effects of new mutations in the local mutational neighborhood of a recently curated, genome-scale metabolic model of *E. coli* (Orth et al., 2011). Our screen included all possible single addition and deletion mutants (Methods), whose growth was simulated on anaerobic glucose media until saturation was reached. Of all non-essential mutations, 147 (3.3%) affected growth rate, either positively or negatively (Fig. 1B). All of these mutations also altered the chemical composition of the environment (see Fig. 1C for a representative subset; Fig. S1 for the full set, Methods), and the magnitude of the environmental and fitness effects were strongly correlated (Pearson’s ρ = 0.61, P<10^-6^, Fig. S2). This suggests that as new mutations fix in the population, the extracellular environment will change, which in turn could alter the fitness effects of new mutations thus deforming the fitness landscape.

We explored the extent of fitness landscape deformability in a dataset that consisted of ~ 10^7^ single and double mutants, representing the entire second-order metabolic mutational neighborhood of *E. coli*. The fitness of each mutant was determined in competition with its immediate ancestor as F_M_= log([X’_M_/X_M_]/[X’_A_/X_A_])(Lenski et al., 1991; Travisano and Lenski, 1996); where X_A_ and X_M_ represent initial densities of ancestor and mutant, and X‘_A_ and X’_M_ their final densities after 10 hours of competition, respectively (see Methods). All competitions were performed at an initial mutant frequency of 0.01. Using this measure, the fitness of two mutations is expected to combine additively when they act independently (Fig 1D). As shown in Fig. 1E, when two mutations without an environmental effect interact with one another, epistasis (ε) will cause the fitness of the double mutant to deviate from additivity. This is the usual definition of epistasis in the literature, and epistasis is, as usual, invariant with respect to the order in which mutations occur (Poelwijk et al., 2007). In contrast, when at least one of the single mutants has an environmental effect, the double mutant experiences a different extracellular environment depending on which of the two single mutants is its immediate ancestor. For example, a double mutant could crossfeed on one of its possible single-mutant ancestors, but not on the other (Fig. 1F). The result is a gene-by-environment-by-gene (G×E×G) interaction in which the magnitude of epistasis depends on the order in which mutations occur. In other words, epistasis becomes non-commutative. The value of that non-commutative fitness shift (δ) characterizes the deformation of a two-step mutational trajectory(Fig. 1F).

### Deformability in the path to an evolutionary innovation in *E. coli*

To experimentally validate this concept and assess the potential relevance of landscape deformability in experimental evolution, we studied two mutations on the path to the evolutionary innovation of strong aerobic growth on citrate (Cit^++^) in the Ara-3 population of the *E. coli* Long-Term Evolution Experiment (LTEE)(Blount et al., 2008). The two principal mutations underlying this phenotype are known to have profound ecological consequences, suggesting that non-commutative epistasis may be present (Fig. 2A). The first mutation is a tandem amplification overlapping the citrate fermentation operon, *cit*, which occurred after 31,000 generations. This amplification caused aerobic expression of the CitT transporter, producing a weak citrate growth phenotype (Cit^+^)(Blount et al., 2012). CitT is an antiporter that imports citrate, which is present in large amounts in the LTEE’s DM25 growth medium, while exporting intracellular C_4_-dicarboxylate TCA intermediates, including succinate and malate (Quandt et al., 2015), thereby increasing their concentration in the extracellular environment. A subsequent mutation causes high level, constitutive expression of DctA, a proton-driven dicarboxylic acid transporter. This mutation refines the Cit^+^ trait to Cit^++^ by allowing recovery of the C_4_-dicarboxylates released into the medium by both the progenitor and the double mutant itself during growth on citrate (Quandt et al., 2014) (Fig. 2A). We reasoned that these mutations together enable the exploitation of environments built by progenitor strains, producing a stronger increase in fitness than expected in the absence of environmental construction (Fig. 2B). In contrast, had the DctA mutation occurred prior to the CitT-activating duplication, it would have conferred no fitness benefit, and would not have produced any changes in the environment relative to the ancestral strain (Fig. 2B).

**Fig. 2.**
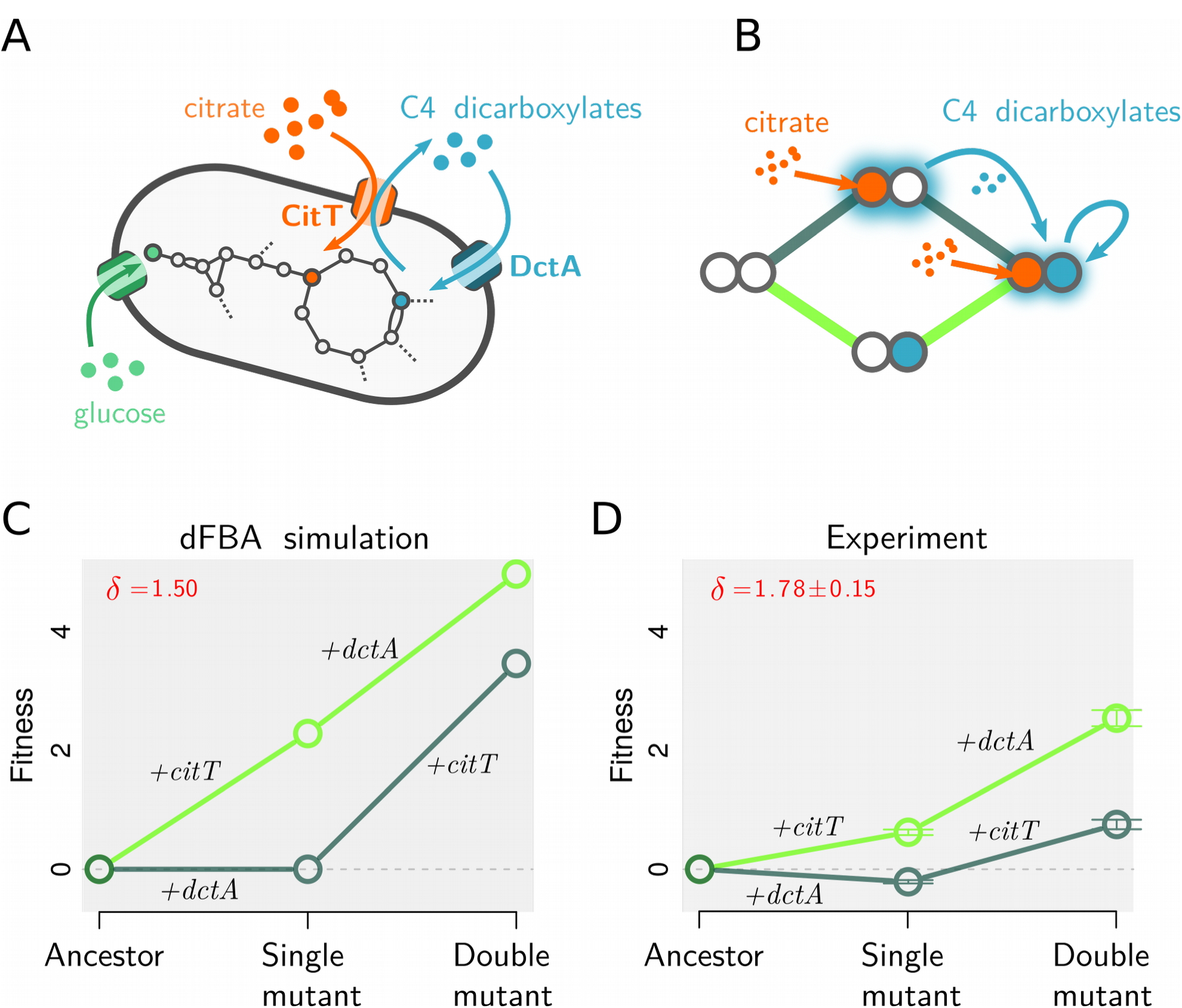
Non-commutative epistasis in the evolution of aerobic citrate use in *E. coli*. **A.** Function of the two transporters involved in the innovation of strong aerobic growth on citrate (Cit^++^) in *E. coli*. CitT is an antiporter that exchanges extracellular citrate for internal C_4_-dicarboxylates (e.g. succinate, fumarate, malate). DctA is a carboxylic acid transporter that imports C_4_-dicarboxylates from the extracellular space into the cytoplasm. **B.** The two possible mutational trajectories leading to the Cit^++^ trait. If the mutation leading to expression of *citT* (+*citT*) occurs first, it will transform the environment leading to cross-feeding towards the double mutant. This should not occur if the *dctA* overexpression mutation (+*dctA*) occurs first. Simulated (**C**) and experimentally measured (**D**) fitness landscapes in the DM25 medium used in the LTEE (see Methods). Experimentally obtained values are reported as mean ± SEM (N=10).

We tested this prediction by performing competitive fitness assays with different combinations of a spontaneous Cit^−^ mutant and *dctA*^−^ knockout strains derived from ZDB89, a 35,000 generation Cit^++^ clone that possesses both the DctA-activating and CitT-activating mutations (see Methods). Competitions were carried out with equal volumes of each combination of competitors, and relative fitness determined using colony counts obtained after 0 and 24 hours of growth (Lenski et al., 1991). In parallel, we used dynamic FBA modeling to simulate these competitions, relying solely on known parameters from the experiments, as well as published parameters pertaining to the physiology of *E. coli* (Gallet et al., 2017; Harcombe et al., 2014)(see Methods). Confirming our expectations, dynamic FBA predicts strong non-commutative epistasis (δ =1.50) (Fig. 2C). This is confirmed by the experimental results (δ=1.78±0.15) (Fig. 2D). The agreement between the empirically calibrated computational model and the experiments is not only qualitative but quantitative: with no fitting parameters, dynamic FBA is predictive of the outcome of the experimental pairwise competitions, explaining 52% of the variance in colony counts from all experiments (N=120; Fig. S3).

### Short-range deformability in *E. coli* is weak and rare

Although the above examples demonstrate the potential presence of fitness landscape deformability, its pervasiveness in empirical fitness landscapes remains unclear. To shed light onto this question, we screened the entire second order mutational neighborhood of *E. coli* using our computational model (Fig. 3A). In Fig. 3B we represent all pairs of mutations that exhibit deformability as nodes in a network that are connected if their non-commutative fitness shift (δ) is larger than 1% of the fitness effects (F_MAX_). These represent only a small subset (203/3343, or 6.1%) of all epistatic interactions, which for the most part are not altered by the environmental effects of mutations.

**Fig. 3.**
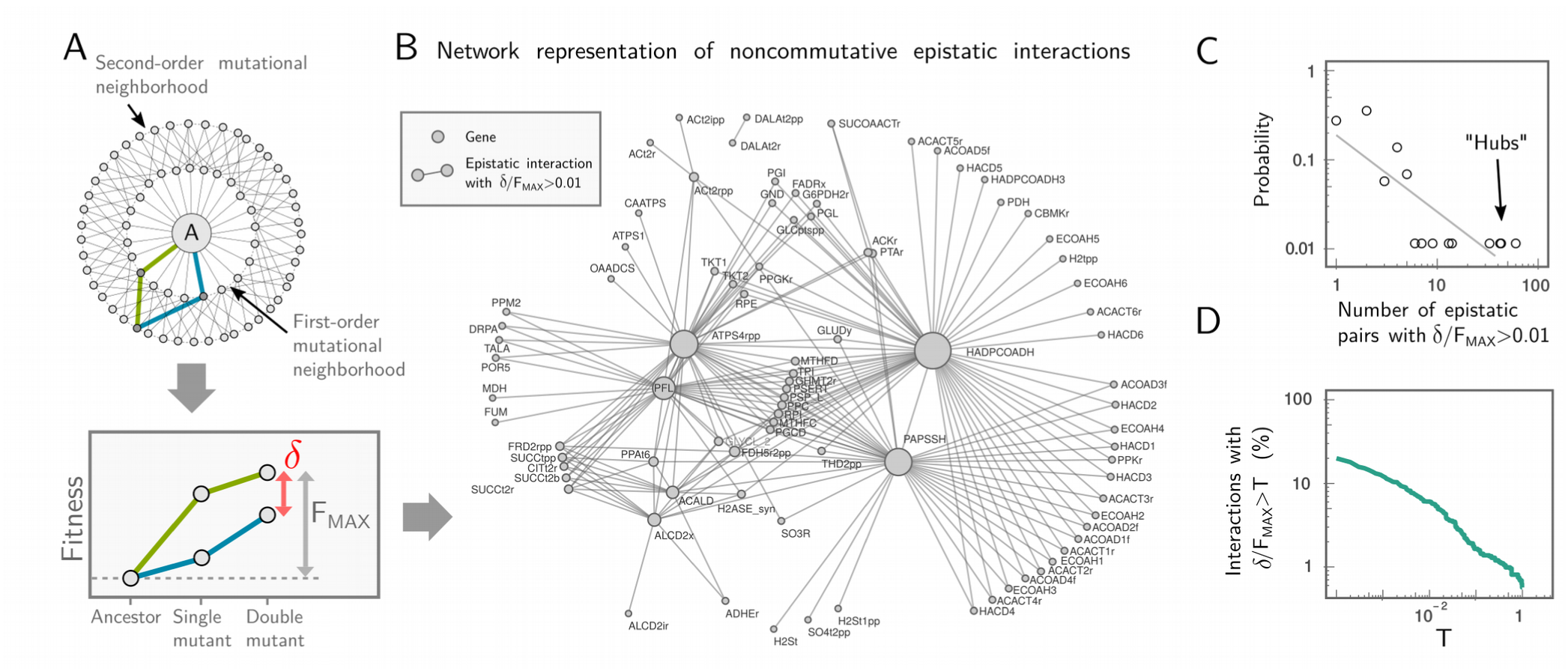
Short-range deformability in *E. coli* is rare, weak, and directional. **A** Systematic exploration of the second-degree mutational neighborhood of *E. coli*. We used our model to exhaustively simulate every possible mutational trajectories starting from the “ancestral”(A) metabolism and ending in each double-mutant, including all nonessential reaction deletions and additions. Non-commutative epistasis (δ) was measured for each pair of mutants, and normalized by F_MAX_, i.e. the maximal cumulative fitness effect of the double mutant: F_MAX_ = max[|F_i_^(A)^+ F_ij_^(i)^|, |F_j_^(A)^ +F_ij_^(j)^|], where e.g. F_x_^(y)^ denotes the fitness of mutant *x* when invading its immediate ancestor *y* at low frequency (see also Methods for simulation details). **B.** Network representation of all non-commutative epistatic pairs. Nodes represent single mutations, and two nodes are joined by edges if δ/F_MAX_ > 0.01 for that epistatic pair. Node names represent reaction names from BIGG database. **C.** Distribution of deformability for each gene in the network, measured as the number of other genes with which it has a non-commutative epistatic interaction. **D.** Strength of all non-commutative epistatic interactions. Percentage of epistatic pairs with δ/F_MAX_ >T as a function of T.

Non-commutative interactions also tend to be unevenly distributed: most of the mutations do not deform the fitness of any other mutation, and only 15 of them (0.3% of all), deform the fitness of 5 or more other mutations (Fig. 3B-C). These few highly connected “hubs” on the network tend to be the mutations with the strongest environmental effects (Fig. S4; Pearson’s correlation = 0.79, P<10^-6^). Non-commutative epistatic effects also tend to be small in magnitude (Fig. 3D); only 1.6% (55/3343) of epistatic pairs have a non-commutative epistatic shift larger than 10% of the total fitness increase (δ/F_MAX_ >0.1, Fig. 3D). This reveals that deformability of the local mutational neighborhood of the *E. coli* metabolic landscape is both generally weak, rare, and highly anisotropic (i.e. non homogenous), with deformations limited to localized directions in genotype space.

### Long-range deformability of the *E. coli* metabolic fitness landscape

The low deformability of the local mutational neighborhood could be explained by the strong genetic similarity between the mutants and the ancestral genotype: genotypically close descendants will rarely be able to use metabolites that are discarded by their immediate ancestors. By the same logic, one may predict that over longer mutational distances metabolic differences might accumulate that enable the use of extracellular metabolites that are left as a “legacy” by previous mutations. Thus, we hypothesize that changes to the extracellular environment produced by a given mutation will primarily deform the fitness landscape at distant positions on the genotype space.

To test this hypothesis, we set out to introduce a mutation with a strong environmental effect and measure the deformation it causes at different distances in the genotype space. We chose the ACKr mutation (the deletion of the acetate kinase gene), which as shown in Fig. 1C modifies the environment by releasing large amounts of lactate at the expense of lower secretions of formate, acetate and ethanol. To quantify the deformation introduced by this mutation, we compared the fitness of thousands genotypes at increasing mutational distances from the ancestor, in competition with either the ancestor *E. coli* model (A) or the ACKr mutant (M) (Fig. 4A). The results are shown in Fig. 4B-C. Consistent with our hypothesis, we found that the fitness landscape deformation introduced by ACKr is negligible at short genotypic distances from it (e.g. 16 mutations or less), but it becomes stronger at longer distances. Fifteen other mutants (M) in addition to ACKr were also tested, with similar results (Fig. S5). Furthermore, by comparing the growth rate of thousands of genotypes in the environments constructed by A and M, we found that increasingly distant genotypes become increasingly sensitive to the differences between both environments. This explains the observed pattern of deformation as a function of genotypic distance (Fig. 4D, Fig S5).

**Fig. 4.**
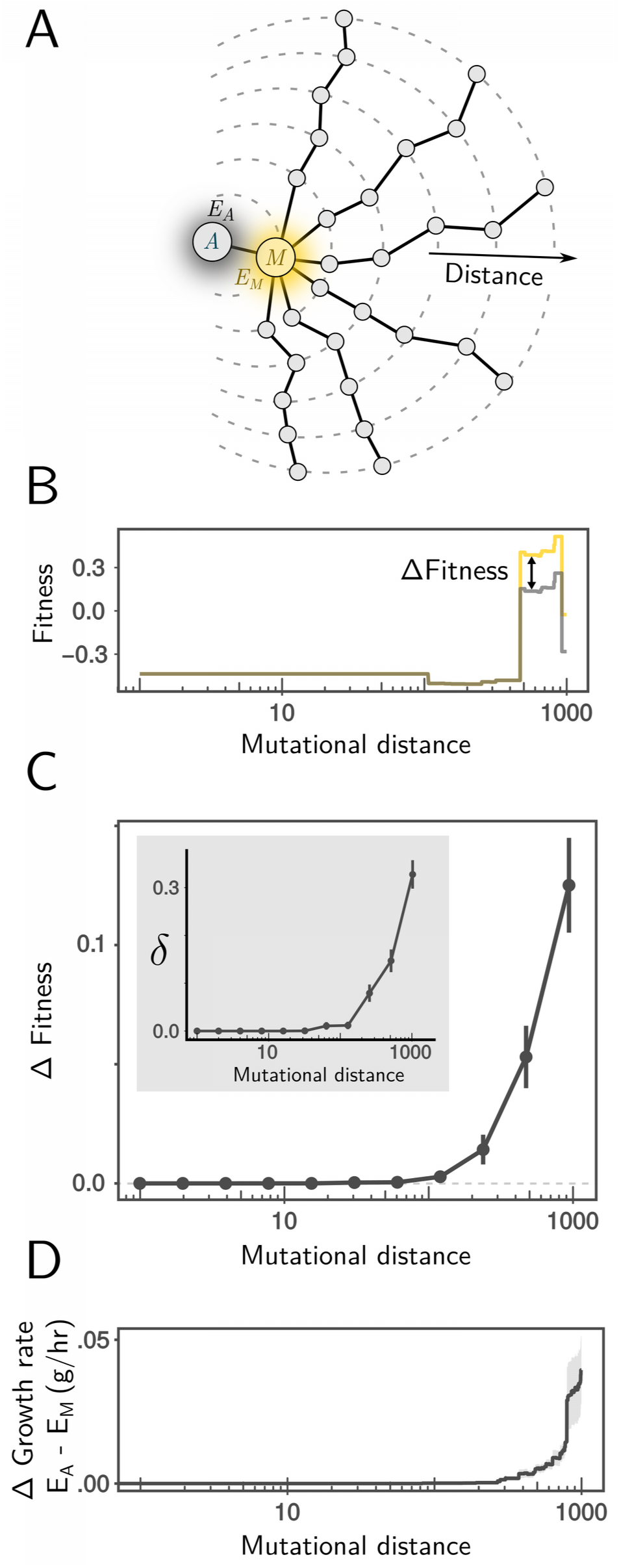
Long-range deformability of the *E. coli* metabolic fitness landscape. **A.** We performed random walks (length = 1000 mutations) in genotype space starting from a *E. coli* ancestor (A; gray) and first passing through a mutant (M; orange) with large environmental effect. **B-C.** Fitness of mutants along these random walk was measured in competition with A (gray), and in the environment it generates (E_A_), as well as in competition with M (orange) in the environment it generates (E_M_). In **B** we show the result for a single example random walk. Note that fitness in competition with M is shifted by the difference in fitness between M and A so all observed differences in fitness (ΔFitness) are due to deformation. In **C** we plot the average ΔFitness at increasing mutational distances from A in over N=100 random walks (error bars represent SEM; N=100). The non-commutative epistasis δ exhibits a similar trend as ΔFitness (inset). **D.** Average difference (absolute value) in growth rate between environments E_M_ and E_A_(in grams of dry cell weight × hr^-1^) at varying genotype distances (gray line, shading represents SEM; N=100).

## Discussion

Darwin was perhaps the first to recognize that the environment experienced by a population can also be shaped by the population itself (Darwin, 1881). Long neglected, this concept was revived by Lewontin (Lewontin, 1978, 1983), and has gained added momentum in recent years as the important role played by eco-evolutionary feedbacks in both ecology and evolution has become better appreciated <u>(Laland et al., 2014; Post and Palkovacs, 2009; Rudman et al., 2018)</u>. Due to technical limitations, experimental studies of eco-evolutionary feedbacks and the adaptive dynamics models that seek to explain them often lack explicit, genome-wide representations of the adaptive landscape, in particular with regard to complex traits and gene-gene interactions (Rudman et al., 2018). The exact state of the environment, which is intrinsically complex and multi-dimensional (Lawrence et al., 2012), is also rarely measured experimentally or explicitly included in eco-evolutionary models. In return, and in spite of early abstract models of species coevolution, which introduced the idea of fitness landscape deformability (also referred to as “rubberness”, ((Kauffman and Johnsen, 1991; Solé and Sardanyés, 2014)), and the many examples of the importance in co-evolutionary arms races and other forms of coevolution (Morran et al., 2011; Stern and Sorek, 2011; Strauss et al., 2005), genotype-fitness maps have largely ignored the effects of eco-evolutionary feedbacks. This is particularly important in light of the argument, made by many authors, that the deformability of fitness landscapes (or its consequences, in the form of frequency dependent selection) would erode their practical and conceptual utility (Doebeli et al., 2017; Moran, 1964; Schuster, 2012).

The work presented above seeks to reconcile both perspectives on empirical grounds. Encouragingly, our results show that fitness landscapes may retain their local properties even in the presence of mutations that significantly alter the environment. By systematically mapping an empirical fitness landscape, we have found that ignoring deformability and assuming a “rigid” (i.e. non-deformable) landscape is a good approximation over short genotypic distances. This is because closely related genotypes are unlikely to differ from one another in their physiological response to the built environment. In contrast, over longer mutational distances, landscapes are likely to be affected by environmental construction, an effect that is shaped by complex genetic interactions. This suggests a new, ecologically-mediated mechanism by which historical contingency may shape downstream evolution even in clonal populations. In summary, our work suggests that depending on the resolution, fitness landscapes can behave as either a fixed externally determined topography on which adaptation proceeds, or become a dynamic property of the populations adapting on them (Doebeli et al., 2017; Moran, 1964).

Our results indicate that simulating cellular adaptive dynamics with an explicit and biologically realistic representation of the genotype-phenotype map is within reach. Such an approach will shed light into the role played by dynamic niche construction on cellular evolution. We believe that it will also create multiple opportunities to incorporate genomics into the study of eco-evolutionary dynamics, and thus reveal the genetic, biochemical and environmental constraints that simultaneously govern the ecology and evolution of cellular populations.

## Acknowledgments

The authors wish to thank Gunter Wagner, Steve Stearns, David Post and members of the Sanchez laboratory for helpful discussions and feedback on the manuscript; and Daniel Segre and Ilija Dukovski for helpful advice and discussions during the early implementation of COMETS. We would also like to thank Maia Rowles, Kiyana Weatherspoon and Brooke Sommerfeld for their assistances in construction of Ara-3-derived strains, and Richard Lenski for use of LTEE strains and materials. We also want to thank Claus Jonathan Fritzemeier and Martin Lercher for assistance with model building and curation. This work was partially funded by a Young Investigator Award (RGY0077/2016) from the Human Frontier Science Program to AS, and a John Templeton Foundation Foundational Questions in Evolutionary Biology grant (FQEB #RFP-12-13) to ZB. The LTEE is supported in part by the National Science Foundation (NSF; DEB-1019989).

## METHODS

### Reconstruction of a prokaryotic genotype space

All *in silico* explorations of genotype space in this work took as a reference the *E. coli* model iJO1366 and consisted of both gene additions or deletions. Gene deletions were performed by constraining both upper and lower bounds of the reaction to zero. Gene additions were performed from a set of all known prokaryotic reactions. We used the BiGG database (King et al., 2016) to compile a dataset of all known reactions found across prokaryotic species. Conflicts in reaction directionality were resolved as follows i) if a reactions is found in the well benchmark *E. coli* iJO1366 model, use the properties given by this model, ii) if a reaction conflicts in directionality, only accept directions found across all models (e.g. if there is one model where a given reaction is irreversible, we set it as irreversible). We used this dataset to create a “universal” metabolic model that included all reactions found in *E. coli* iJO1366 as well as a set of all potential novel reactions. We removed reactions that would lead to erroneous energy-generating cycles using the ModelFit algorithm (Fritzemeier et al., 2017). The algorithm was constrained to conserve reactions present in the original *E. coli* model. Removing any futile cycles from this “universal” model ensures that there will not be any futile cycles in any subset. The resulting network contains 4999 metabolic reactions and 585 nutrient uptake or sink reactions, of which 2758 and 255 were not found in the original *E. coli* model.

### *In silico* simulation of growth through metabolic modeling

Dynamic Flux Balance Analysis simulations were performed using the COMETS package (“Computation of Microbial Ecosystems in Time and Space”, (Harcombe et al., 2014)) and the gurobi optimizer software. For computationally intensive simulations, we used the High Performance Facility at Yale University. For standard (non dynamic) FBA simulations, we used the COBRApy python package (Ebrahim et al., 2013). Both Dynamic and Standard FBA optimizations were done using the parsimonious algorithm, in which a first optimization is done to maximize biomass yield, and a second one fixes this yield and minimizes total fluxes throughout the network (Lewis et al., 2010). Unless otherwise stated, the default V_max_ was set in dynamic FBA simulations to 10 mmol×gr^-1^×hr^-1^ for all uptake reactions. Inorganic ions and gases where kept at high concentrations and where kept undepleted throughout the simulation (i.e lower bound: -1000 mmol×gr^-1^×hr^-1^, amount of metabolite: 1000 mmol).This was done to constrain our analysis to situations where growth is limited only by uptake of carbon sources. The unbounded nutrients are: ca2_e, cbl1_e, cl_e, co2_e, cobalt2_e, cu2_e, fe2_e, fe3_e, h_e, h2o_e, k_e, mg2_e, mn2_e, mobd_e, na1_e, nh4_e, ni2_e, pi_e, sel_e, slnt_e, so4_e, tungs_e, zn2_e. For the citrate simulation to avoid oxygen, nitrogen or proton limitation uptake was unconstrained by setting the vmax to 1000 mmol×gr^-1^×hr^-1^. Analysis of results was performed using GNU R language (R Core Team, 2017).

### Fitness, environmental effects and deformability measurements

To measure fitness, we use here (in both experiments and simulations) the Malthusian fitness measure that allows for a quantitative comparison across environments (Wagner, 2010). Fitness of mutant M relative to ancestor A is therefore given as F_M_= log([X’_M_/X_M_]/[X’_A_/X_A_]), where X and X’ represent initial and final densities. For a pair of mutations, deformability can be then measured as δ_ij_ = F_ij_^(i)^ + F_i_^(A)^-(F_ij_^(j)^+ F_j_^(A)^), where F_x_^(y)^ represents the fitness of genotype *x* in competition with genotype *y*. To compute environmental effects of mutations, the difference in secretion profile of mutants (as shown in Fig. 1C and Fig. S1) is computed for a given released molecule as sign(D) * (log(D)+1) where D is the amount released by the mutant minus that secreted by Ancestral *E. coli*. This log-modulus transformation (John and Draper, 1980) is applied to help visualization of the generally small differences in released amount, which can be either positive or negative. To measure environmental effect of a mutation (as used in Fig. S2 and Fig. S4), we use the Euclidean distance in the profile of released metabolic byproducts between a mutant and the *E. coli* ancestor using standard Flux Balance Analysis (Ebrahim et al., 2013).

### Simulation of the fitness landscape of *E.coli* citrate utilization

Starting with *E.coli* model iJO1366 we constructed metabolic models of the four mutants necessary to predict the fitness landscape involved in the evolution of aerobic citrate utilization in the Ara3 population of the LTEE. Unlike the LTEE ancestral strain REL606 (and *E. coli* generally), which possess the necessary genes for citrate utilization but do not express them in aerobic conditions, iJO1366 is able utilize both citrate and succinate if these reactions are unbounded (as FBA optimizes precisely regulation). Thus, the ancestral phenotype was recreated by knocking out three reactions CITt7pp (*citT*), SUCCt2_2pp (*dctA*) and SUCCt2_3pp (*dcuA* or *dcuB*). The reactions encoded by the first two genes (*citT* and *dctA*) are known to be involved in the evolution of citrate utilization in the LTEE whereas *dcuA* and *dcuB* are involved in dicarboxylate uptake in anaerobic conditions and are inactive in aerobiosis (Six et al., 1994). This triple knockout represents the pre-citrate *E. coli* ancestor strain. The addition of CITt7pp simulates the promoter capture and consequent aerobic expression of CitT. Similarly, the addition of SUCCt2_2pp is equivalent to the first mutation (aerobic expression of *dctA*). We used dynamic FBA to predict the fitness landscape of these two mutations, calibrating the simulations to reflect the the experimental conditions. This involved i) setting the *in silico* media to reflect DM25 minimal glucose media (0.139mM glucose, 1.7mM citrate). Aerobic condition was simulated by keeping oxygen (o2_e) undepleted. ii) using published parameters pertaining to the physiology of *E. coli* (Harcombe et al., 2014) and iii) estimating the initial biomasses of each mutant prior to competition. Initial biomass for citrate simulations was determined using initial plate counts from pairwise competitions experiments (see also Fig. S3). We assume that average cell dry mass is 3.9 * 10^-13^ g which is the empirically measured cell dry mass of REL606 the ancestral strain used in the LTEE (Gallet et al., 2017)

### *E. coli* Long-Term Evolution Experiment

Briefly, twelve populations of *E. coli* B were founded in 1988 from clone REL606. The populations were initially identical, save for half having a mutation that permitted growth on arabinose. (See below.) These have since been evolved in DM25 minimal glucose medium under conditions of daily, 100-fold serial transfer, and incubation at 37°C with 120 rpm orbital shaking. Samples of each population are frozen every 500 generations ^38^. DM25 is Davis-Mingioli broth supplemented with 25 mg/L glucose. (Per liter: 7g potassium phosphate dibasic trihydrate, 2g potassium phosphate monobasic anhydrous, 1g ammonium sulfate, 0.5g sodium citrate, 0.01% magnesium sulfate, and 0.01% thiamine.)

### Isolation and Preparation of Test Strains

ZDB89 is a Cit^++^ clone isolated from the Ara-3 population sample frozen for generation 35,000 during the LTEE. Cit^−^ revertants arise spontaneously from Cit^+^ and Cit^++^ clones due to recombination-mediated collapse of the tandem cit amplification to the ancestral genotype at that locus. We isolated a Cit^−^ revertant, ZDB757, by first passaging ZDB89 in a glucose-only medium for five days. This passaging does not constitute a selection, but nonetheless enriches for Cit^−^ revertants by eliminating the selective penalty for losing the ability to grow on citrate. Passage cultures were spread on LB plates, and Cit^−^ mutants screened for by patching colonies to LB and Minimal Citrate (MC) plates to identify clones that no longer grew on citrate. The Cit^−^ phenotype was confirmed by streaking on Christensen’s Citrate Agar. Recombineering with the pKO3 suicide plasmid (Link et al., 1997) was used to delete the *dctA* gene from ZDB89 and ZDB757, producing the Cit^+^ *dctA*^−^ and Cit^−^ *dctA*^−^ constructs, ZDB912 and ZDB904, respectively. To permit differentiation of competitors during fitness assays, we isolated Ara^+^ revertants of each of the aforementioned clones and constructs. Briefly, Ara^−^ strains lack the ability to use arabinose, and form red colonies on Tetrazolium Arabinose (TA) plates, while Ara^+^ revertants are mutants with restored ability to grow on arabinose, and form white colonies on TA. The ancestral strain of the Ara-3 population and its descendants are Ara^−^. We isolated Ara^+^ revertants by plating clone or construct cultures on Minimal Arabinose (MA) plates. Revertants were competed against their Ara^−^ parents to verify marker state neutrality. Clones, constructs, and revertants are listed in Supplementary Table 1. Derivation of constructs and revertants are shown in Supplementary Figure S7.

### Experimental fitness Assays

Fitness was assayed in pairwise competitions. Competitors with opposite Ara marker states were inoculated from frozen stocks into 10 mL LB broth, and incubated overnight at 37°C with 120 rpm orbital shaking to permit revival and elimination of traces of glycerol cryoprotectant. To precondition the competitors, each competitor revival culture was then diluted 100-fold in 0.85% saline, and 100 L of the diluted culture used to inoculate 9.9 mL DM25 with ten-fold replication. These culture were grown for 24 hours at 37°C with 120 rpm orbital shaking, after which they were transferred via 100-fold dilution into 9.9 mL volumes of fresh DM25, and grown for another 24 hours under the same conditions. Ten competition cultures were prepared for each competitor pairing by inoculating each 9.9 mL DM25 with 50 L of each preconditioned competitor. A single replicate preconditioning culture of each competitor for each competition was inoculated so that each competition was inoculated from a single preconditioned culture of the competitors. Upon inoculation with the competitors, 100 L of a 100-fold dilution of each was spread on TA to permit enumeration of the initial frequency of each competitor. 100 L of a 1000-fold dilution was also plated for each competition including at least one Cit^+^ or Cit^++^ competitor. Colonies were counted following 48 hours of plate incubation at 37°C. Following 24 hours incubation under the same conditions used for preconditioning, 100 L of 10,000-fold dilutions of each competition were plated on TA to permit final enumeration of the competitors. 100,000-fold dilutions were also plated for competitions including at least one Cit^+^ or Cit^++^ competitor.

### Exploration of deformability in the local mutational neighborhood of *E. coli.*

To systematically analyze the local mutational neighborhood of *E.coli* we construct a set of metabolic models consisting of every viable single and double mutation, considering both additions and deletions from our universal reaction set and using as a reference the *E. coli* iJO1366 model (4389 and 9636050 genotypes, respectively). We removed from the final analysis essential genes, as well as those genes leading to artifacts (H_2_ or CO_2_ limitation). We used dynamic flux balance analysis to simulate competition assays of each mutant with its immediate ancestor. The simulations assayed co-culture growth during 10hr, a period during which glucose was never exhausted, i.e. growth remained exponential, to simplify the interpretation of the results. All simulations were done with the mutant starting at low frequency (1%, 10^-10^ gr. dry cell weight, for 9.9 × 10^-9^ gr. for the ancestor) in anaerobic glucose minimal media (unless otherwise stated, see detailed parameters in supplement and supplementary tables ST2-ST4).

### Simulation of long-range fitness landscape deformation

In order to explore the long-range effects of landscape deformation, we started from a one-step mutation from ancestor *coli* model and performed random walks in genotype space by sampling 1024 mutations (without replacement) among both deletions and additions. To speed-up simulations, the sampling procedure ignored all reactions that were essential in a minimal model built by sequentially removing reactions while possible, following (Pál et al., 2006)). At regular intervals, fitness was measured as before in competition with the mutant and the ancestor (wild-type) using dynamic flux balance analysis (COMETS (Harcombe et al., 2014)). To determine the growth rates of genotype in ancestral vs mutant environments we repeated this procedure except at each step, growth rate was measured in the environment provided by the mutant and the ancestor using standard flux balance analysis (COBRApy (Ebrahim et al., 2013)). These environments were simulated by setting uptake rates for each secreted metabolites to the excretion rate of the respective ancestor.

